# Cell cycle re-entry in the aging *Drosophila* brain

**DOI:** 10.1101/2024.08.26.609689

**Authors:** Deena Damschroder, Jenny Sun, Katherine O. McDonald, Laura Buttitta

## Abstract

The brain is an organ comprised mostly of long-lived, quiescent cells that perform vital functions throughout an animal’s life. Due to the brain’s limited regenerative ability, these long-lived cells must engage unique mechanisms to cope with accumulated damage over time. We have shown that a subset of differentiated neuronal and glial cells in the fruit fly brain become polyploid during adulthood. Cell cycle re-entry in the brain has previously been associated with neurodegeneration, but there may be a more complex relationship between polyploidy and cell fitness in the brain. Here, we examine how known lifespan modifiers influence the accumulation of polyploidy in the aging fly brain. Flies aged at a low temperature, or with a low protein diet, accumulate polyploid cells in the brain more slowly than expected if this phenotype were solely regulated by lifespan mechanisms. Despite the slower accumulation of polyploid cells, animals under conditions that extend lifespan eventually reach similar levels of polyploidy in the brain as controls. Our work suggests known lifespan modifiers can influence the timing of cell cycle re-entry in the adult brain, indicating there is a flexible window of cell cycle plasticity in the aging brain.

## Introduction

Cells in the brain accumulate DNA damage during aging which can lead to cell death when too severe (Fishel et al., 2007; Folch et al., 2012; Kruman, 2004). Consequently, cell death triggered from age related DNA damage contributes to neurodegeneration (Folch et al., 2012). This is problematic because the brain is comprised mostly of cells thought to be in a permanent, non-proliferative state of G0 (Aranda-Anzaldo, 2012; Aranda-Anzaldo & Dent, 2017). Therefore, other mechanisms besides proliferation may compensate for cell loss in the aging brain.

Cells in G0 are metabolically active and carry out various physiological functions but are not actively cycling. As research in the cell cycle field has progressed, flexibility in the G0 state has been realized, even for terminally differentiated cells in vivo (Borowik et al., 2023; Losick, 2016; Ma et al., 2019; Nandakumar et al., 2021; Yao, 2014; Zanet et al., 2010). In some cases, terminally differentiated cells can re-enter the cell cycle and undergo variant cell cycles to compensate for tissue damage, demonstrating surprising plasticity in the cell cycle and G0 state (Cohen et al., 2018; Dehn et al., 2023).

The flexibility of G0 in differentiated cells has not been extensively examined in the brain. It is generally believed that cells in an adult brain, especially neurons, are in a permanent G0 state (Folch et al., 2012; Kruman, 2004). Despite being in a stable non-proliferative state, multiple studies report cells with a more than 2C DNA content (polyploid cells) in the brain and variant cell cycle gene reactivation (Jungas et al., 2020; Khurana & Feany, 2007; Lu et al., 2004; McCarroll et al., 2004; Sigl-Glockner & Brecht, 2017; Sosunov et al., 2020; Yang & Herrup, 2007). This suggests cells in the brain, including neurons, thought to be in a stable G0 state, retain the ability to re-enter the cell cycle. Consistent with this, we have shown that under normal, physiological conditions, the adult fly brain accumulates a small proportion of polyploid cells, likely due to cell cycle re-entry in adults (Nandakumar et al., 2020). The presence of polyploid terminally differentiated cells in the aged brain under normal and diseased conditions indicates an intriguing, but poorly understood cell cycle plasticity in the brain.

The evidence for quiescent cells in the adult *Drosophila* brain that re-enter the cell cycle is relatively sparse, and there are many unknowns, including how polyploid neurons and glia might arise. Neuro-glial progenitors have been reported in the optic lobes (Fernandez-Hernandez et al., 2013; Simoes et al., 2022), central brain (Crocker et al., 2021; Kato et al., 2009), and antennal lobes (Fernandez-Hernandez et al., 2021; von Trotha et al., 2009), that engage DNA replication when the adult brain is injured (Foo et al., 2017; Kato et al., 2009; Li et al., 2020; von Trotha et al., 2009). Evidence for mitotic divisions exists but they appear to be rare, suggesting daughters of progenitors could undergo mitotic slippage or mitotic skipping to result in polyploid cells. Alternatively, differentiated diploid neurons or glia may re-enter a variant cell cycle lacking mitosis in response to tissue damage or cell loss, in a manner similar to wound-induced polyploidization, observed in the adult Drosophila epithelium (Besen-McNally et al., 2021; Grendler et al., 2019; Losick, 2016).

Regardless of whether the source of polyploid cells in the adult brain is neuro-glial progenitors, differentiated cells, or a combination of both, the very low levels of polyploidy in brains of newly eclosed adults indicates most cell cycle re-entry from a G0 state occurs during the first few weeks of adulthood. We previously showed that polyploid cells accumulate most rapidly during the first three weeks of adulthood, reaching a plateau by 21 days of age (Nandakumar et al., 2020). The steady state of polyploidy after 3 weeks may be due to reduced cell cycle re-entry, or continued cell cycle re-entry balanced by the loss of polyploid cells through latent reductive cell divisions or cell death, or a combination of both. The window of polyploid cell accumulation therefore is an age-associated trait, that can be explored to decipher factors influencing cell cycle plasticity in the brain.

Here, we show that known lifespan modifiers can influence the timing and rate of the accumulation of polyploid cells in the adult fly brain. Manipulations that extend lifespan slow the accumulation of polyploid cells in the brain, but in a manner that cannot be explained by the prolonged lifespan alone. When we scale polyploid cell accumulation to the percentage of maximum lifespan under each condition, normalizing for lifespan extension does not fully resolve the slowed dynamics of polyploid cell accumulation. Collectively, our data suggests there is a relationship between low temperature responses, protein intake and the window of flexibility of G0 in the adult brain. We propose that polyploidy in the brain is the result of a highly regulated process controlled by multiple physiological inputs that results in a reproducible proportion of polyploid cells in the brain to be achieved and maintained during aging.

## Materials and Methods

### Fly Strains and husbandry

The *Drosophila melanogaster w^1118^* fly strain (BDSC 5905) was used for all experiments. Virgin female and male flies collected within 48 hours of eclosion were considered age matched and placed in separate vials containing 20 flies per vial. Flies were housed in a 25°C incubator on Bloomington Cornmeal food and exposed to 12-hours of ambient light per day unless otherwise noted.

### Flow Cytometry

The proportion of polyploid cells in whole brains was determined using flow cytometry. For region specific experiments the optic lobes (OLs) and central brain (CB) from the brain were manually separated. Unless otherwise stated, one female brain and one male brain was used per biological replicate. 3-6 biological replicates were performed per group. DNA content was measured using the live DNA stain, DyeCycle violet. Cell death was measured using permeability assays with either propidium iodide (PI) or Sytox Green. The flow rate was 500 µl per minute for each sample and a minimum of 10,000 events were gated as non-doublets. Flow cytometry experiments were performed as previously described (Nandakumar et al., 2020) on an Attune NxT flow cytometer.

Briefly, fly brains were dissected and dissociated using trypsin:EDTA and then stained with DyeCycle violet and PI or 7-AAD. Brains were incubated for twenty minutes in the dark and then triturated using a p200 pipette tip. The solution was moved to an Eppendorf tube containing 400 µl of the trypsin:EDTA, PBS, and dye solution. The sample was incubated at room temperature for 45 minutes in the dark. After incubation, the sample was diluted with 500 µl of 1X PBS and gently vortexed prior to running on the Attune NxT flow cytometer for analysis.

To identify polyploid cells in our sample we used a stringent gating strategy that was used previously (Nandakumar et al., 2020). Debris was eliminated based on size (FSC) and complexity (SSC). We identified cells by gating around events that were positively stained with DyeCycle Violet. The DNA-stained events were then gated based on area (VL1-A) and height (VL1-H) four doublet elimination. Next, we analyzed the DNA content of non-doublet, DNA-stained events and examined cell death using PI or 7-AAD positivity.

### Fixing, Immunostaining, and Imaging

*Drosophila* brains were dissected in 1X phosphate buffered saline (PBS). Per condition, 4 brains were dissected within 20 minutes to minimize tissue damage. Tissues were fixed in 4% paraformaldehyde (PFA) in 1X PBS for 30 minutes. Tissues were permeabilized in 1X PBS+ 0.05% Triton X-100 and blocked with 1X PBS, 1% BSA, 0.1% Triton X-100 (PAT). Lamin (DSHB ALD67.10) (1:500 dilution) staining was performed in PAT solution overnight at 4°C with rotation. After the overnight incubation, the samples were washed, blocked with PBT-X (1X PBS, 2% Normal goat serum, 0.03% Triton X-100) and incubated with secondary antibody overnight at 4°C. After the secondary antibody incubation, samples were washed with 1X PBS and DAPI was applied for ten minutes. Afterwards, the samples were washed again with 1X PBS and then mounted in Vectashield H1000 (Vector Labs).

All images were taken on a Leica SP8 at 63x magnification with 0.5 micron Z-sections. The OLs and the CB were imaged at age day 7, day 14, and age day 21. Image quantifications were performed as previously described (Box et al., 2024; Grendler et al., 2019). Raw integrated intensity measurements were taken from diploid cells, which comprise most of the brain. Polyploid cells were identified based on nuclear size and circularity of the Lamin stain. The raw integrated intensity was measured, and the background fluorescence was subtracted from each measurement to ascertain the corrected fluorescence intensity. The intensity of haploid DNA content was calculated by dividing the average intensity of the diploid cells by 2. Polyploid cells identified were then confirmed based on the following binning: 2N (1.9–2.9), 4N (3.0–6.9), 8N (7–12.9). Four brains per time point were examined, with 4 OLs images and 2 CB images being analyzed from each brain. Data from all brain regions is pooled together for each timepoint.

### Hydroxyurea Feeding

10mg/ml Hydroxyurea (HU) in 50% sucrose was used to block DNA synthesis in adult brains (Elkahlah et al., 2020). Briefly, HU (ThermoFisher, AAA1083106) was dissolved in 50% sucrose to create a stock of 50 mg/ml. 600 µl of the stock solution was added to reheated Bloomington Cornmeal food for a total volume of 3 ml for a final concentration of 10 mg/ml. Food was made one day prior to the addition of flies. *w^1118^* flies were collected the day they eclosed. Virgin female and male flies were placed on HU or vehicle containing food. Vials were flipped every other day and flow cytometry measurements were taken at age day 7, day 14, and age day 21.

For larvae experiments, yeast paste containing 5 mg/ml of HU dissolved in ddH_2_0 was used instead of solid food (Elkahlah et al., 2020). *w^1118^* flies laid eggs on grape agar plates the day before for two hours. 24-hours post egg legging, the agar plates were checked hourly. Any larvae that emerged within the hour were immediately moved to yeast paste containing 5mg/ml of HU. Larvae were left on the yeast paste for eight hours and then moved to regular cornmeal food. The vehicle group was treated in a similar way, accept the yeast paste contained no HU, only ddH_2_0. For the control group, eggs were laid on the standard Bloomington Cornmeal food. Flow cytometry experiments were performed on adult brains exposed to HU on age day 7.

### Lifespan assays

Female and male *w^1118^* flies were collected over a 24-hour period and separated into different vials to prevent mating. 13 vials of males and 13 vials of female flies containing 20 flies of the same sex per vial were collected per condition. A total of at least 230 flies per condition were measured. Conditions included 25°C, 18°C, 29°C, reduced yeast protein (RY), 25°C + RY, 18°C + RY, and 29°C +RY. Lifespan measurements were performed by changing the food every Monday, Wednesday, and Friday with the number of dead flies recorded each day (Damschroder et al., 2018; Linford et al., 2013). Flies were housed based on the appropriate condition in the dark, which does not influence polyploid cell accumulation (Nandakumar et al., 2020). Two separate cohorts for each condition were examined in parallel. Flies from the different cohorts had different parents. Significance was tested by a log-rank test, or one-way ANOVA of the average median and maximum lifespan.

### Measuring polyploidy under different conditions and scaling to total lifespan

Flies were housed in the various conditions described above. Polyploidy measurements started on the day of eclosion (age day 0), day 7, and then every two days thereafter. Data was scaled to the total lifespan for each condition. The percent lifespan was determined by taking the age in days of the measurement divided by the maximum lifespan of that group then multiplied by 100 (% lifespan = (age in day / maximum lifespan) * 100).

### Graphs and Statistics

Graphs and statistical analysis were created and performed using GraphPad Prism 10. Analysis for all experiments is reported in the figure legends.

## Results

### Resolving the window for cell cycle re-entry in the adult *Drosophila* brain

Incidences of cell cycle re-entry have been reported in *Drosophila* adult brains of mutants and disease or damage models (Crocker et al., 2021; Fernandez-Hernandez et al., 2013; Khurana et al., 2006; Simoes et al., 2022). Cell cycle re-entry also occurs in healthy adult fly brains but is less understood (Foo et al., 2017; Kato et al., 2009; Li et al., 2020; Nandakumar et al., 2020; von Trotha et al., 2009). We previously identified polyploid neuronal and glial cells in the adult *Drosophila* brain under normal physiological aging conditions without mating (Nandakumar et al., 2020). Here we determined whether there is an adult-specific window of cell cycle re-entry in the *Drosophila* brain under these normal physiological conditions.

The proportion of cells containing more than a diploid (2C) DNA content (polyploid cells) increases in female and male wild-type flies during normal physiological aging at 25°C on standard cornmeal and agar food (Fig. 1A, 2-way ANOVA, Aging factor, p<0.0001). The proportion of polyploid cells in the whole brain is similar between female and male flies (Fig. 1A, 2-way ANOVA, Sex factor, p=0.931), and peaked on average at 21-28 days of age (Fig. 1A. Sidak’s multiple comparison tests, p>0.05), which may indicate there is a physiological requirement for a precise proportion of polyploidy in the brain. The proportion of cells with a more than 4C DNA content (greater than tetraploid) is similar in female and male fly brains during aging and peaked on average at 28 days of age (Fig. 1B., 2-way ANOVA, Aging factor, p<0.0001, Sex factor, p=0.652). We measured cell death in the brain using a propidium iodide (PI) cell permeability assessment. The fraction of cells incorporating PI increases during aging with an increase after 21 days of age (Fig. 1C., 2-way ANOVA, Aging factor, p<0.0001) with similar proportions in female and male flies (Fig. 1C., Sex factor, p=0.652). This level of cell death in the adult brain is similar to the cell death rates measured using other permeability assays such as 7-AAD (SF. 1A.).

**Figure 1:**
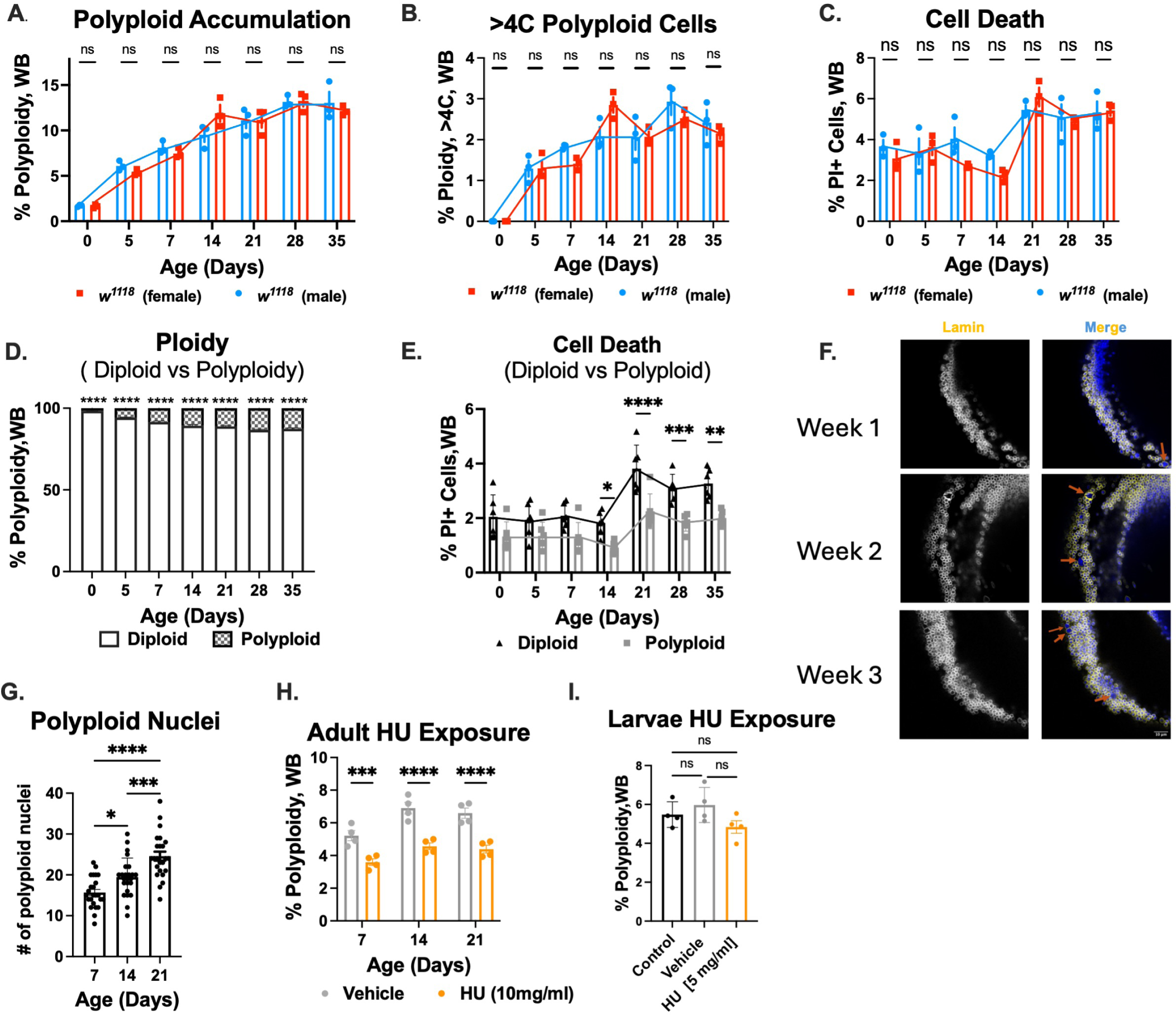
Cell cycle plasticity in the adult fly brain. (A.) Adult female and male flies accumulate polyploid cells similarly during aging (3 biological replicates per time point, 2 fly brains per biological replicate) (2-way ANOVA, Aging factor, p<0.0001, Sex factor, p=0.931). (B.) The proportion of cells that contain more than a 4C (tetraploid) DNA content changes with age similarly in female and male flies (2-way ANOVA, Aging factor, p<0.0001, Sex factor, p=0.652). (C.) Cell death is not significantly different in female and male aging brains (2-way ANOVA, Sex factor, p=0.159), but does increase with age (2-way ANOVA, Aging factor, p<0.0001). (D.) The brain remains mostly diploid during aging (Pooled data, 2-way ANOVA, Ploidy factor, p<0.0001, Sidak’s multiple comparisons test). (E.) Diploid cell death increases with age (2-way ANOVA, Aging factor, p<0.0001, Ploidy factor, p<0.0001, Sidak’s multiple comparisons test) (F.) Representative images of the *Drosophila* optic lobe (OL) (63x magnification) at age day 7, day 14, and age day 21. The orange arrows indicate enlarged, polyploid cells. (G.) Dapi intensity measurements from fluorescent images of the OL and central brain (CB) at day 7, day 14, and day 21 show an increase in polyploid nuclei with age (n=4 brains per timepoint with 4 OL images and 2 CB images per brain) (One-way ANOVA, p<0.0001). (H.) Flies fed the S-phase inhibitor hydroxyurea (HU) from the day of eclosion accumulate fewer polyploid cells during aging (3 biological replicates per time point containing one male and one female brain per biological replicate) (2-way ANOVA, Drug factor, p<0. 0001, Aging factor p=0.003, Tukey’s multiple comparisons test). (I.). Feeding HU to larvae did not prevent polyploidy in adult brains age day 7 (3 biological replicates per aging point, 2 fly brains per biological replicate) (One-way ANOVA, p=0.154). Graphs display individual biological replicates, with SEM. *p<.05, **p<.01 \*\*\**p*<0.001, ***p<0.0001, ns, not significant.

After observing no sex-specific effects on polyploidy or cell death, one female and one male brain were combined in a biological replicate for all subsequent experiments. Even though polyploidy in the brain increases with age, most cells in the brain remain diploid at all time points (Fig. 1D., 2-way ANOVA, Ploidy effect, p<0.0001). Thus, only a small proportion of cells re-enter the cell cycle in adult brains under normal physiological aging conditions. The majority of dead and dying cells in the brain are also diploid (Fig. 1E., 2-way ANOVA, Ploidy effect, p=0.0006), with the proportion of dying diploid and polyploid cells increasing with age (Fig. 1E., 2-way ANOVA, Aging effect, p<0.0001). Consistent with our previous observations, the proportion of dying cells that are polyploid is less than diploid cells at several timepoints (Fig. 1E.) (Nandakumar et al., 2020). Differences in cell death rates between diploid and polyploid cells may be explained by the fact that diploid cells comprise most of the brain.

We verified our flow cytometry results by performing fixed imaging and DAPI intensity measurements on brains at days 7, 14, and 21. These measurements were done similarly to prior studies, except that we used a nuclear lamin staining to manually segment individual nuclei (Besen-McNally et al., 2021; Box et al., 2024) in the optic lobes and central brain using confocal microscopy. Most cells in the brain are diploid and uniform in nuclear circularity and size, while polyploid cells are visually distinguishable with enlarged nuclei, increased DAPI intensity and decreased nuclear circularity (Fig. 1D.). Integrated DAPI intensity measurements from similar-sized z-stacks of the optic lobe and central brain reveal an increase in the number of polyploid nuclei with age, consistent with our flow cytometry data (Fig. 1G. one-way ANOVA, p<0.0001).

On the day of eclosion (day 0), less than 2% of cells are polyploid in the brain, but by day 21 around 10% of cells in the brain are polyploid. Polyploidy can arise through cell fusion, as well as a variant cell cycle known as an endocycle, which includes S-phase (DNA synthesis phase) but lacks mitosis and cytokinesis resulting in increased nuclear ploidy (Nandakumar et al., 2021). To confirm that cell cycle re-entry resulting in endocycling gives rise to polyploid cells in the adult fly brain, we fed adult flies the DNA synthesis inhibitor hydroxyurea (HU) to block S-phases (Koc et al., 2004). Flies fed HU from the day of eclosion develop significantly fewer polyploid cells than vehicle-fed flies (Fig. 1H. 2-way ANOVA, Drug effect p<0.0001) and do not continue to accumulate polyploid cells during aging (Fig. 1H., Tukey’s multiple comparisons test, p>0.05). Therefore, reducing DNA synthesis limits polyploidy in the aging brain and supports the hypothesis that some polyploid accumulation relies on cells re-entering the cell cycle to replicate their DNA.

To ensure the cell cycle- re-entry window is adult specific, we fed HU to larvae for the first 8 hours of life. Flies fed HU as larvae develop adult-specific polyploidy normally during their first week of life (Fig. 1I., One-way ANOVA, p=0.154), demonstrating that blocking DNA synthesis before adulthood does not influence polyploid cell accumulation. In addition, this demonstrates that cell types in the brain depleted by larval HU feeding such as the mushroom body neuroblasts in the central brain, are not a significant source of polyploid cells in the adult brain. While we cannot rule out the possibility that neural or glial progenitor cells observed by others (Fernandez-Hernandez et al., 2013; Kato et al., 2009; von Trotha et al., 2009) contribute to the accumulation of polyploidy in the brain, these cells are believed to be largely quiescent in the adult brain under normal physiological conditions (Fernandez-Hernandez et al., 2013; Ito & Hotta, 1992; Kato et al., 2009; Li et al., 2020; Li & Hidalgo, 2020; Siegrist et al., 2010; Truman & Bate, 1988; von Trotha et al., 2009). Our data therefore supports a model where some of the age-associated accumulation of polyploid cells under normal physiological conditions is due to cells re-entering the cell cycle during the first 3 weeks of adulthood.

### Extending lifespan delays cell cycle re-entry in the adult brain

Due to the association between aging and the accumulation of polyploidy in the brain, we next examined whether known lifespan modifiers could influence the timing or rate of polyploid cell accumulation. Housing temperature and protein content in diet are known lifespan modifiers (Grandison et al., 2009; Mair et al., 2005; Molon et al., 2020; Shaposhnikov et al., 2022; Skorupa et al., 2008; Tatar et al., 2014). We first confirmed the effects of altered housing temperatures on lifespan, specifically a low (18°C) and a high (29°C) temperature. Consistent with previous reports (Miquel et al., 1976; Shaposhnikov et al., 2022) female and male flies housed in 18°C on the day of eclosion (age day 0) live longer than their age-matched siblings housed at the control temperature (25°C) (Fig. 2A.), while flies housed at 29°C have a significantly shorter lifespan than their age-matched siblings housed at 25°C (Fig. 2B.).

**Figure 2:**
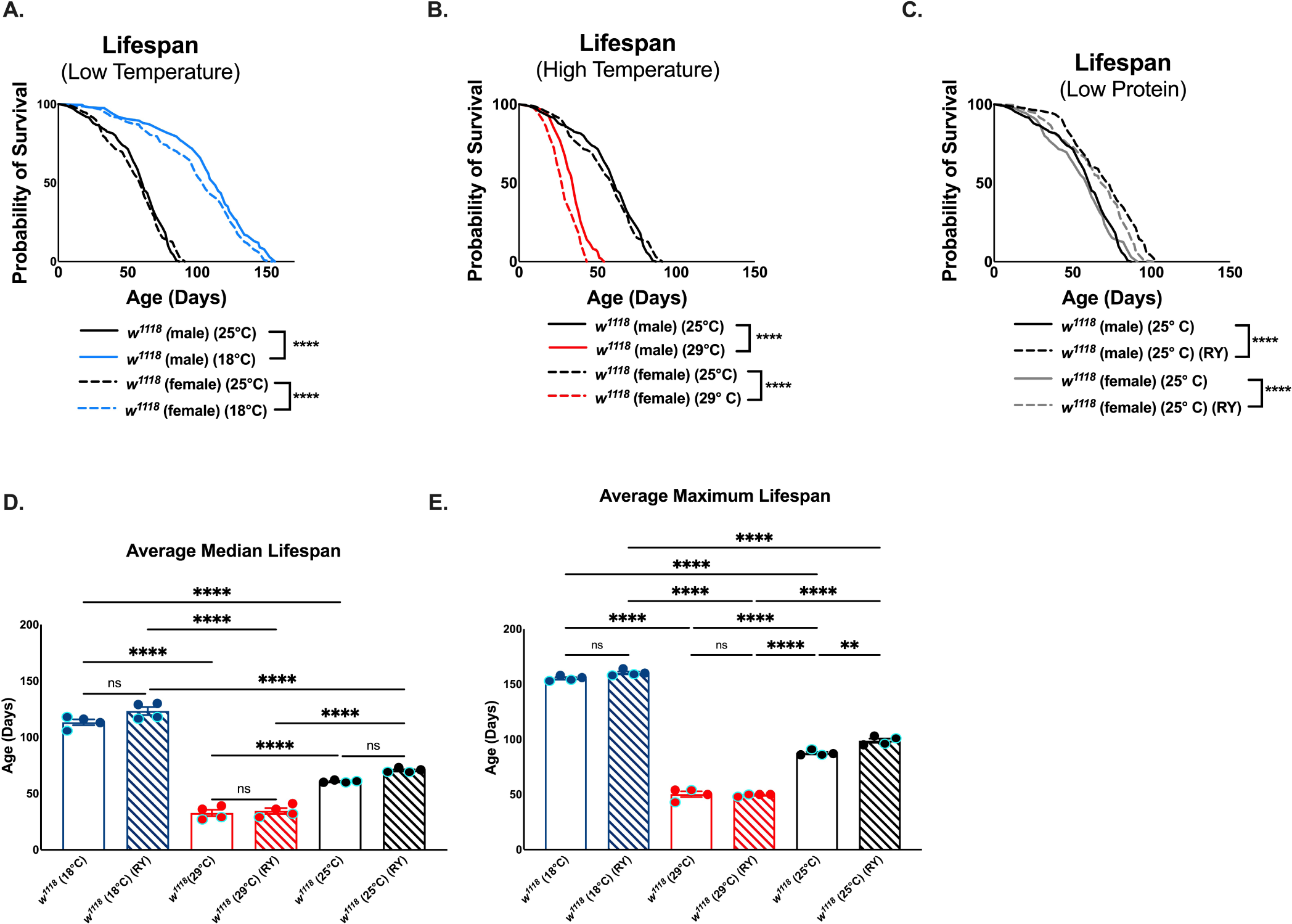
Temperature and protein levels in diet influence adult fly lifespan. (A.-C.) One replicate of lifespan assessment under different housing conditions. 260 flies were measured for each condition and lifespan curves were compared, using a pair-wise log-rank analysis. Significant differences are noted (\*\*\*\**p*<0.0001). (A.) Flies housed at 18°C live longer than their sex-matched siblings (male (25°C) vs male (18°C), p<0.0001, female (25°C) vs female (18°C), p<0.0001). (B.) Flies housed at 29°C live shorter than their sex-matched siblings (male (25°C) vs male (29°C), p<0.0001, female (25°C) vs female (29°C), p<0.0001). (C.) Reduced yeast (RY) diet increases lifespan similarly in males and females (male 25°C vs female 25°C, p=0.725, male 25°C RY vs female 25°C RY, p= 0.322, RY vs. control diet p<0.0001). (D. & E.) The average median and maximum lifespan from 2 replicate lifespan assessments were calculated (one-way ANOVA with Tukey’s multiple comparisons test). Graphs display the average from all replicates with SEM. Replicates from adult females are outlined in turquoise. Each replicate contains 260 flies per condition. The effects of temperature on lifespan are much greater than RY diet. RY diet did not significantly increase average lifespan at any temperature, but did increase maximum lifespan at 25°C.

We next confirmed the effects of feeding a reduced protein diet by restricting yeast content, (Grandison et al., 2009; Steck et al., 2018) which is the main source of amino acids and protein in the standard cornmeal and agar fly food (https://bdsc.indiana.edu/information/recipes/bloomfood.html). Our reduced yeast (RY) diet contains 30% less yeast than control food (1.1% total yeast in RY compared to 1.58% yeast in control diet). Feeding rate decreases with age (SF2) (Min & Tatar, 2006) and there is a small, but significant, increase in feeding rate in young flies fed a reduced yeast diet (SF 2A., 2-way ANOVA, Diet effect, p=0.020). At age day 21 there is no difference in feeding rate (SF 2B., 2-way ANOVA, Diet effect, p=0.80). Flies fed the RY diet from age day 0 live longer than their age-matched siblings fed the control diet (Fig. 2C.), although this lifespan extension is subtle, which may be explained by the fact that our control diet is already a lower yeast diet (1.58%) compared to other common diets (Ren et al., 2020; Skorupa et al., 2008).

When replicate lifespans were combined and averaged, flies aged at a low temperature had the largest average median and maximum lifespan (Fig. 2D. & E.) while flies aged at a higher temperature had the lowest average median and maximum lifespan (Fig. 2D & E.), as expected based on prior work (Miquel et al., 1976; Shaposhnikov et al., 2022; Zajitschek, 2023). The effects from combining modifiers on lifespan is unclear, with some groups finding additive effects (Phillips, 2024; Shaposhnikov et al., 2022) while others do not (Zajitschek, 2023). We observed no additive effects to the average median and maximum lifespan when combining these different lifespan modifiers. For example, housing flies at 18°C while on a low protein diet has no effect on median (Fig. 2D. Tukey’s multiple comparison test, p=0.083) or maximum lifespan (Fig. 2E., Tukey’s multiple comparison test, p=0.272), compared to 18°C on our normal diet. Furthermore, there is no rescue to the median (Fig. 2D. Tukey’s multiple comparison test, p=0.995) or the maximum (Fig. 2E., Tukey’s multiple comparison test, p=0.993) lifespan in flies housed at 29°C and fed a low protein diet. Nevertheless, altered temperature and RY diet alone were reproducible lifespan modifiers.

We then examined the effects of these lifespan modifiers on polyploid cell accumulation in the adult brain. Flies housed at 18°C accumulate polyploid cells in their brain significantly slower than flies housed at 25°C (Fig. 3A., 2-way ANOVA, Temperature factor, p<0.0001) and the proportion of cells with a more than 4C DNA content is also reduced (Fig. 3B. 2-way ANOVA, Temperature factor, p<0.0001). Cell death in flies housed at 18°C is not significantly altered (Fig. 3C., 2-way ANOVA, Temperature factor, p=0.173), suggesting the changes in polyploid cell accumulation at 18°C cannot be explained by differences in cell death in the brain. By contrast, flies housed at 29°C accumulate polyploidy similarly to flies at 25°C (Fig. 3D., 2-way ANOVA, Temperature factor, p=0.641). However, the proportion of cells containing more than 4C DNA content is partially influenced by aging at a high temperature (Fig. 3E., 2-way ANOVA, interaction, p=0.0003), with flies at 29°C exhibiting more cells with higher (>4C) DNA content by day 35 (Fig. 3E., Sidak’s multiple comparison test, p=0.0002). This increase in cells with higher ploidies is not due to significant changes in cell death however (Fig. 3F. Temperature factor, p=0.914). This suggests there could be a relationship between the stress of aging at a high temperature and developing cells of higher ploidies than tetraploid at later timepoints in adulthood.

**Figure 3:**
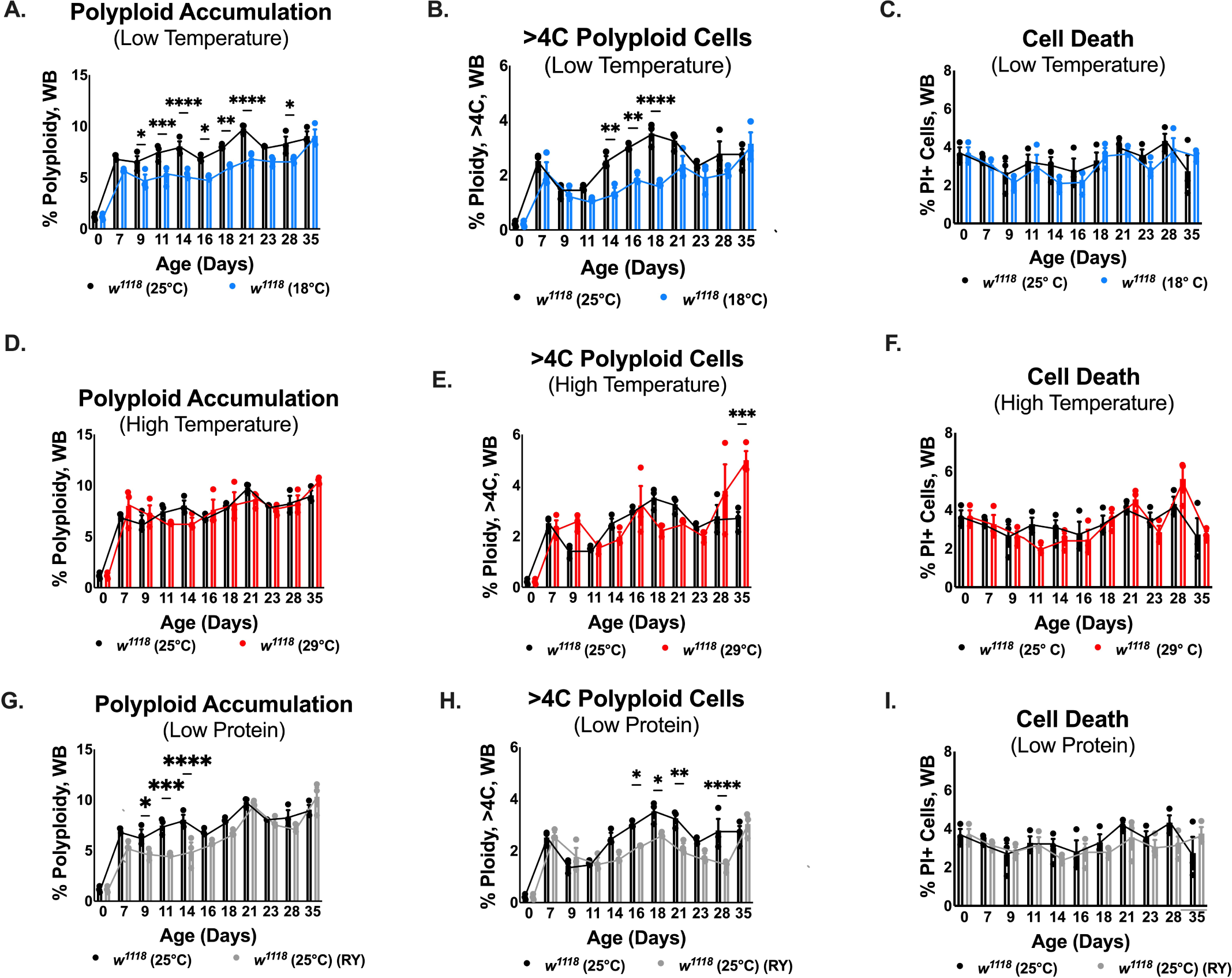
Conditions that increase lifespan delay cell cycle re-entry in the adult fly brain. (A.) Flies housed at 18°C accumulate polyploid cells in their brain slower than flies housed at 25°C (2-way ANOVA, Aging factor, p<0.0001, Temperature factor, p<0.0001, Sidak’s multiple comparisons test). (B.) Flies housed at 18°C accumulate fewer cells with a DNA content exceeding 4C (2-way ANOVA, Aging factor, p<0.0001, Temperature factor, p<0.0001, Sidak’s multiple comparisons test). (C.) Cell death is not significantly influenced by housing temperature (2-way ANOVA, Aging factor, p=0.001, Temperature factor, p=0.173,). (D.) Flies aged at 29°C show no significant differences in polyploid cell accumulation in the brain compared to flies aged at 25°C (2-way ANOVA, Aging factor, p<0.0001, Temperature factor, p=0.641). (E.) However the proportion of cells with a DNA content exceeding 4C significantly increased with age at 29°C (2-way ANOVA, Aging factor, p<0.0001, Interaction, p=0.0003, Sidak’s multiple comparisons test). (F.) Housing flies at 29°C did not significantly impact cell death in the adult brain (2-way ANOVA, Aging factor, p<0.0001, Temperature factor, p=0.914). (G.) Flies fed a low protein diet (30% reduced yeast, RY) exhibit slowed accumulation of polyploid cells in the brain (2-way ANOVA, Aging factor, p<0.0001, Diet factor, p<0.0001, Sidak’s multiple comparisons test). (H.) Low protein-fed flies accumulate fewer cells with a DNA content exceeding 4C during aging (2-way ANOVA, Aging factor, p<0.0001, Diet factor, p<0.0001, Sidak’s multiple comparisons test). (I.) Cell death was not influenced by the low protein diet (2-way ANOVA, Diet factor, p=0.171). Graphs display individual biological replicates, with SEM. All significant differences are noted: *p<.05, **p<.01 \*\*\**p*<0.001, ***p<0.0001.

Like reduced temperature, flies on a low protein diet (RY) accumulate polyploidy slower in the brain (Fig. 2G., 2-way ANOVA, Diet factor, p<0.0001) and fewer cells with a DNA content of more than 4C (Fig. 2H., 2-way ANOVA, Diet factor, p<0.0001). Slower polyploid accumulation in the brain is not due to altered cell death in flies fed a low protein diet (Fig. 2I., 2-way-ANOVA, Diet factor, p=0.171). In fact, the proportion of polyploid cell death in the whole brain is significantly reduced in flies fed a reduced protein diet (SF3 E., 2-way ANOVA, Diet factor, p<0.0001). The slower accumulation of polyploid cells during the first 2 weeks of adulthood may contribute to the reduced number of cells with higher ploidies more than 4C at later ages, beyond 2 weeks (Fig. 3G, H.).

Combining lifespan modifiers indicate an interaction between a high housing temperature and protein availability. Flies housed at 18°C and fed a low protein diet and flies housed at 29°C and fed a low protein diet had less polyploid cell death than controls (SF. 3G., 2-way ANOVA, Temperature and diet effect, p=0.0314, SF 3. I, 2-way ANOVA, Temperature and diet effect, p=0.001).

Flies housed at 18°C and fed a low protein diet accumulate the same proportion of cells that contained a more than 4C DNA content as flies housed at 18°C (Fig. 4B., 2-way ANOVA, Temperature and Diet factor, p=0.0056,). Therefore, combining a low protein diet with a low housing temperature does not further reduce the development of cells with higher ploidies. In contrast, when we compared the proportion of cells with a more than 4C DNA content at 29°C and 29°C with a low-protein RY diet, we observe a significant reduction of higher ploidies at 4 weeks of age (Fig. 4E., 2-way ANOVA, Temperature & Diet factor, p=0.007).

**Figure 4:**
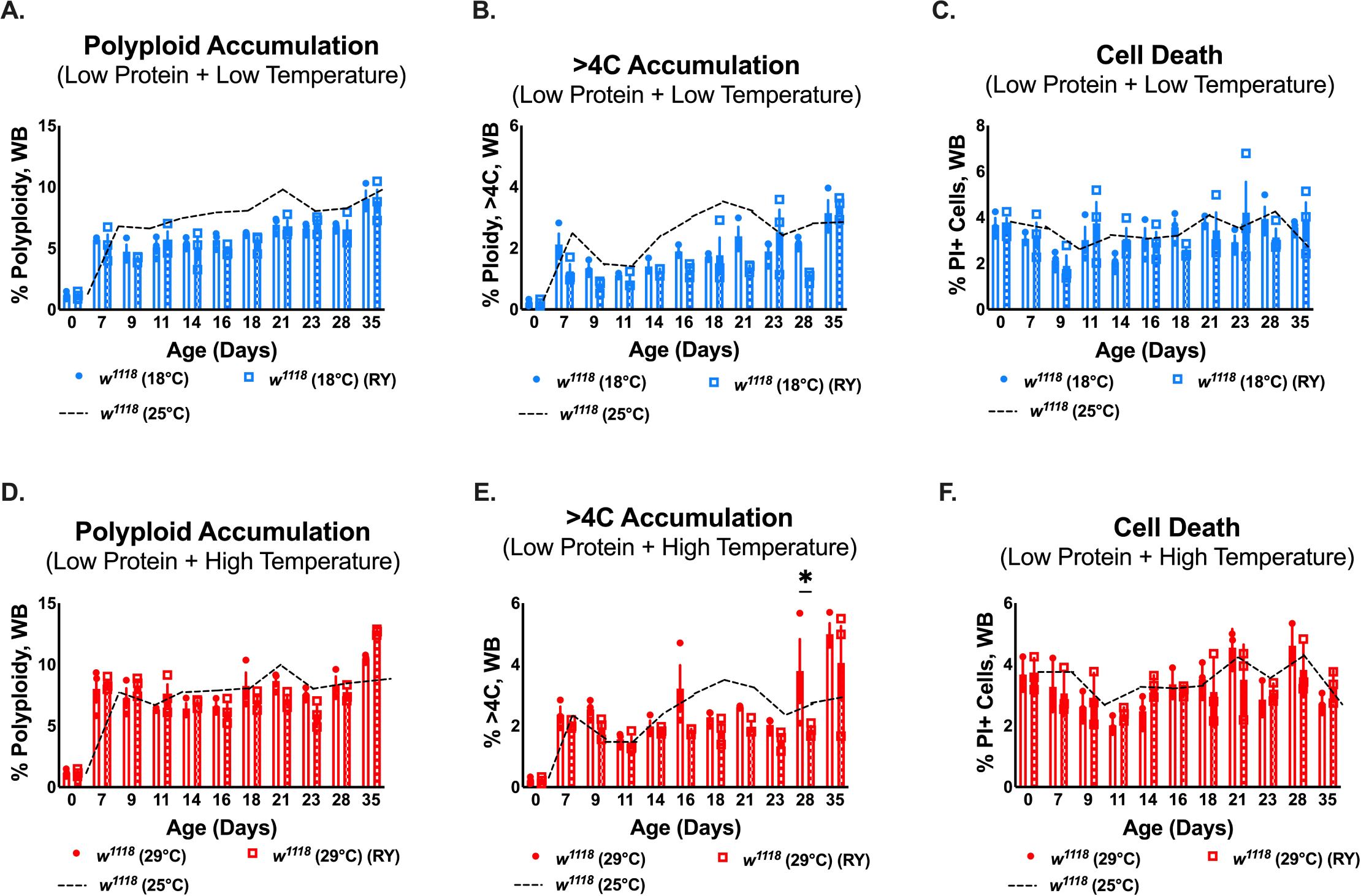
The accumulation of cells with a higher than 4C DNA content may rely on protein availability. (A.) overall polyploidy in the brain is similar between flies housed at 18°C and flies housed at 18°C and fed a low-protein diet (2-way ANOVA, Diet & Temperature factor, p=0.423). (B.) Combining a low temperature and a low protein diet mildly increases the accumulation of cells with a DNA content that is more than 4C relative to just housing at low temperature (2-way ANOVA, Aging factor, p<0.0001, Temperature & Diet factor, p=0.0056, interaction, p=0.041, Sidak’s multiple comparisons test). (C.) Housing flies at a low temperature in addition to feeding a reduced protein diet has no impact on cell death (2-way ANOVA, Aging factor, p=0.057, Temperature & Diet factor, p=0.669). (D.) Feeding a low protein diet to flies housed at a high temperature does not influence polyploid accumulation (2-way ANOVA, Aging factor, p<0.0001, Temperature & Diet factor, p=0.879). (E.) However, the number of cells with more than 4C DNA content is influenced by housing flies at a high temperature with reduced protein (2-way ANOVA, Aging factor, p<0.0001, Temperature & Diet factor, p=0.007, Sidak’s multiple comparisons test). (F). Housing flies at a reduced temperature in addition to feeding a reduced protein diet has no significant impact on cell death (2-way ANOVA, Aging factor, p=.057, Temperature & Diet factor, p=0.876). Graphs display individual biological replicates, with SEM. Each biological replicate contains one male and one female brain. All significant differences are noted: *p<.05, **p<.01 \*\*\**p*<0.001, ***p<0.0001.

Flies housed at 29°C exhibit an increased number of cells with higher ploidies (Fig. 3E.), while flies fed a low-protein diet at a normal temperature (25°C) accumulate fewer cells with higher ploidies (Fig. 3H.). Our data suggests that a lack of sufficient protein in the diet hinders the development of highly polyploid cells in the adult brain at higher temperatures, indicating that the cell cycle re-entry process in the adult brain may be partially dependent upon protein and amino acid sensitive signaling pathways.

### The cell cycle re-entry window in the adult brain is flexible

Collectively, our data demonstrates that conditions that extend lifespan delays cell cycle re-entry in the adult brain resulting in the slower accumulation of polyploid cells and reduced levels of cells with higher ploidies. However, despite the slower rate of polyploid cell accumulation, the proportion of polyploid cells in the brain eventually reaches control levels. The underlying reason for this is unclear but implies the proportion of polyploidy in the brain may be physiologically relevant. This also suggests the window of cell cycle re-entry is not dependent upon chronological age but may scale with physiological age. Therefore, we next examined whether the accumulation of polyploid cells under lifespan alterations scaled with percent of maximum lifespan which effectively normalizes the phenotype to physiological age, rather than chronological age (number of days after eclosion).

When we plot polyploid cell accumulation based on percent maximum lifespan, we see that after an initial increase immediately after eclosion, polyploid cell accumulation is still significantly slower under conditions of reduced temperature or reduced protein (RY), even when normalized to the total lifespan under these conditions (Fig. 4B.). If the window of cell cycle re-entry in the adult brain is entirely dependent upon physiological aging cues, we would expect the levels to overlap when normalized to total lifespan. The reduced levels of polyploid cells in the brain at 18°C or on a reduced protein diet suggests that the window for cell cycle re-entry in the adult brain is flexible and responds to signaling impacted by temperature and protein levels in a manner that is more sensitive to, or independent of the downstream pathways responsible for lifespan extension.

### Polyploidy accumulation is brain region specific

Different regions of the *Drosophila* brain accumulate different levels of polyploid cells (Nandakumar et al., 2020). We therefore examined whether polyploidy in different brain regions may be affected differentially by aging at a low temperature or being fed a low protein diet. To examine this, we physically separated optic lobes (OLs) from the central brain (CB) and performed flow cytometry under the indicated lifespan extending conditions to assess whether effects on cell cycle re-entry may be brain region specific. We consistently observe the levels of polyploidy in the OLs are more affected by temperature and protein levels relative to the central brain. The OLs of flies housed at 18°C accumulate significantly fewer polyploid cells (Fig. 6A., 2-way ANOVA, Temperature factor, p<0.0001) and fewer cells of high ploidies (Fig. 6B., 2-way ANOVA, Temperature factor, p=0.0031), than the CB, which showed no significant difference in the accumulation of polyploid cells under these conditions (Fig. 6D. 2-way ANOVA, Temperature factor, p=0.420, Fig. 6E., >4C DNA content, 2-way ANOVA, Temperature factor, p=0.295). Unexpectedly, cell death in the CB is significantly increased at a reduced temperature for a single timepoint (Fig. 6F.) but is not significantly affected in the OLs at any timepoint (Fig. 6C.), suggesting the changes in the levels of polyploid cells of the OLs are not caused by changes in cell survival. The OLs of flies on the low protein diet also accumulate fewer polyploid cells than flies on a control diet (Fig. 7A., interaction p=0.0021). while the CB shows no effect of a low protein diet on polyploid cell accumulation (Fig. 7D. Diet factor, p=0.314). Interestingly, like flies aged at a low temperature, flies fed a low protein diet have significantly more cell death in their CB at one timepoint (Fig. 7F., 2-way ANOVA, Diet factor, p=0.0068). While the reasons why low temperature and low protein diet may affect cell death in the central brain is unclear, this finding again suggests the effects on polyploid cell accumulation in the OL are not indirectly due to changes in cell survival.

Our data supports the possibility that cell cycle re-entry in the adult fly brain and the polyploid cell accumulation that results, is responsive to temperature and amino acid signaling, possibly through metabolic activity, specifically in the OL. This important result separates the effects of these pathways on polyploidy from their effects on lifespan and suggests the cells that respond to these cues to induce cell cycle re-entry are specific to, or enriched in, the optic lobes.

## Discussion

### An adult-specific window for cell cycle re-entry in the brain

We previously found an onset of polyploidization in the adult fly brain (Nandakumar et al., 2020). Here, we resolve the window for cell cycle re-entry in the adult brain, which relies on DNA replication during the first three weeks of adult life. When flies are housed under conditions that extend lifespan, we see the rate of polyploid cell accumulation in the brain is slowed at a rate independent of lifespan extension. Furthermore, we show that the brain region influences polyploid cell accumulation. Collectively, our data points to an adult-specific program in the brain that integrates information from signaling pathways impacted by temperature and protein content in the diet, to establish a window for cell cycle re-entry in the optic lobes.

The mechanistic Target of Rapamycin (mTOR) network, specifically mTOR complex 1 (mTORC1), could be a key regulator for this adult-specific cell cycle re-entry program in the aging brain. mTORC1 integrates nutrient availability, growth factor signals, and cellular energy to induce various pathways that drive cell growth and proliferation (Weichhart, 2018). To achieve increased cell growth and proliferation, mTORC1 activity increases nucleotide synthesis, protein synthesis, and DNA replication licensing through S6K, 4E-BP1, and Cdc6 respectively (He et al., 2021), all of which are also required for cell cycle re-entry. Interestingly, in flies aged at 18°C, mTOR activity is reduced (Scialo et al., 2015), which could explain the slowed rate of polyploid cell accumulation we observed in flies aged at 18°C. Future studies will illuminate if the mTOR signaling network impacts the cell cycle plasticity of the aging brain.

### Some cells in the adult brain are in a flexible state of G0

Work over the past decade supports the non-proliferative G0 state of differentiated cells is not a binary on or off switch, but more of a spectrum with some cells having more flexibility to re-enter the cell cycle, while other cells maybe in a stable, robust G0 (Borowik et al., 2023; Losick, 2016; Ma et al., 2019; Nandakumar et al., 2021; Yao, 2014; Zanet et al., 2010). Our work suggests there is heterogeneity in the flexibility of G0 among cells in the brain and reveals that environmental conditions influence the plasticity of the G0 state.

In this study, we focused on how polyploid cell accumulation changes in the whole brain, OL, or CB under different housing conditions during aging. We see that housing adults at a reduced temperature or feeding a reduced protein diet slows the accumulation of polyploidy in the whole brain. These manipulations do not block cell cycle re-entry, since polyploid cells still accumulate with age. Additionally, feeding the DNA synthesis inhibitor, hydroxyurea, during adulthood significantly reduced the age-associated polyploid cell accumulation (Fig. 1H.). The slower accumulation under low temperature or reduced protein could be due to a reduced number of cells that re-enter the cell cycle, a slower speed of the variant cell cycle, or some combination of these two possibilities. Further work will be needed to resolve these important open questions.

When polyploid cell accumulation is examined by brain region, we see the OLs specifically accumulate polyploid cells slower when housed at a reduced temperature or fed a reduced protein diet. This suggests either the flexibility of the G0 state is brain region specific, or that the cell type(s) poised to re-enter the cell cycle in adult brains are specific to or enriched in the OLs. Several neuronal and glial cell types become polyploid in adults (Nandakumar et al., 2020), so further work will be needed to resolve the sources and lineages of adult-specific polyploid cells in the brain.

### Polyploid cells in the brain may arise from multiple origins and different triggers

The adult fly brain is mostly post-mitotic and does not normally retain developmental neuroblasts (Siegrist et al., 2010). However, after direct physical injury to the brain in either the optic lobe (OL) (Fernandez-Hernandez et al., 2013; Simoes et al., 2022) or the central brain (CB) (Crocker et al., 2021; Kato et al., 2009), cell division and neurogenesis ensue, suggesting quiescent neuro-glial progenitors exist in these regions. BrdU (Kato et al., 2009) and EdU (Foo et al., 2017) labeling shows DNA replication occurring in these same regions in young wild-type brains without injury up to age day 10, but the mitotic marker PH3 is very rarely present (Kato et al., 2009). In our previous work, we also saw no convincing evidence of mitosis in aged wild-type flies (Nandakumar et al., 2020). The positive S-phase labeling and lack of PH3 labeling is indicative of cells entering a non-mitotic variant cell cycle. Currently, it is unclear whether adult neuro-glial progenitor cells are contributing to the physiological age-dependent accumulation of polyploid cells, or whether cell cycle re-entry of terminally differentiated cells occurs. Alternatively, a combination of differentiated and progenitor cells may contribute to polyploidy in different regions of the brain. Further work will require lineage tracing to decipher the potentially complex potential sources of polyploid cells in the adult fly brain. Regardless of the source of polyploid cells, our data indicates there is adult-specific cell cycle re-entry occurring in the brain.

The stimuli that trigger polyploidization in the brain also remain unclear. Acute physical injury to the fly brain (Crocker et al., 2021; Fernandez-Hernandez et al., 2013; Foo et al., 2017), acute DNA damage in the fly brain (Nandakumar et al., 2020), and overexpression of known neurodegeneration-inducing proteins (Khurana et al., 2006; Rimkus et al., 2008) can induce cell cycle activity. In this study, we did not cause any acute physical injury, DNA damage, or express neurodegenerative proteins. Our study specifically examines cell cycle re-entry during normal physiological aging. It is known that DNA damage naturally accumulates in the brains of flies (Fishel et al., 2007; Nandakumar et al., 2020). Relative to acute damage, DNA damage during aging may be slower and more prone to engage adaptive mechanisms. As such, this may explain why the increase in polyploidy in the brain, and by extension cell cycle re-entry, is less dramatic during aging relative to the cell cycle activity that occurs from direct damage or genetic manipulation paradigms.

A possible hypothesis is only specific stressors influence the rate of polyploidization in the brain. For example, housing flies at a higher temperature, which significantly reduces lifespan, does not accelerate the accumulation of polyploidy in the brain but increases the proportion of cells exhibiting higher ploidies (>4C). This suggests that the rate of cell cycle re-entry may be a response to specific triggers while the degree of polyploidization may respond to chronic stress.

### Possible models to explain the accumulation of polyploidy in the aging brain

In control flies, the total proportion of polyploidy in the brain increases with age and then plateaus after age day 21 (Fig 1A.). This suggests the brain obtains a physiological proportion of polyploid cells that is maintained through an unknown mechanism during aging. We establish that reduced temperature or reduced protein slows the initial increase of polyploidy in the brain, in a manner partially independent of the lifespan increase. The mechanisms involved in achieving and maintaining this physiological proportion of polyploid cells in the brain are unresolved.

To maintain the plateau observed (Fig. 1A.), the polyploid cells may continue to be produced but progress through mitotic or amitotic reductive divisions to become diploid in the aged brain. We do not observe an increase in diploid cells in the aging brain (Fig. 1D.), and find little evidence for mitosis in wild-type fly brains (Foo et al., 2017; Kato et al., 2009; Li et al., 2020; Nandakumar et al., 2020; von Trotha et al., 2009). As there are no molecular markers for amitosis (Lucchetta & Ohlstein, 2017) this possibility will be difficult to address and live cell imaging in adult brains will be challenging since polyploid cells accumulate over weeks.

A different possibility is that a subset of cells re-enters the cell cycle, become polyploid, and then no further cell cycle re-entry occurs later in adulthood. This hypothesis suggests that the G0 state becomes more robust with age, which is supported by other studies (Buttitta et al., 2010; Buttitta et al., 2007; Ma et al., 2019). Interestingly, housing at a reduced temperature or feeding a reduced protein diet slows the accumulation of polyploidy in the brain (Fig. 3) but not the overall total (Fig. 5), demonstrating that the window of G0 flexibility is modifiable. Temperature or diet manipulations influence overall metabolism in the fly (Asiimwe et al., 2023; Blanchard et al., 2024; Klepsatel et al., 2016; Klepsatel et al., 2019). Therefore, metabolism and metabolic mediators may be involved in determining the robustness of G0 in early adulthood. A promising mediator between metabolism and the cell cycle re-entry in the brain is the insulin-Akt-mTOR pathway, which is known to influence the endocycle in the fly (Ohhara et al., 2017; Xiang et al., 2017; Zielke et al., 2011). Future work will determine the impact this metabolic pathway and others have on G0 robustness in the aging brain.

**Figure 5:**
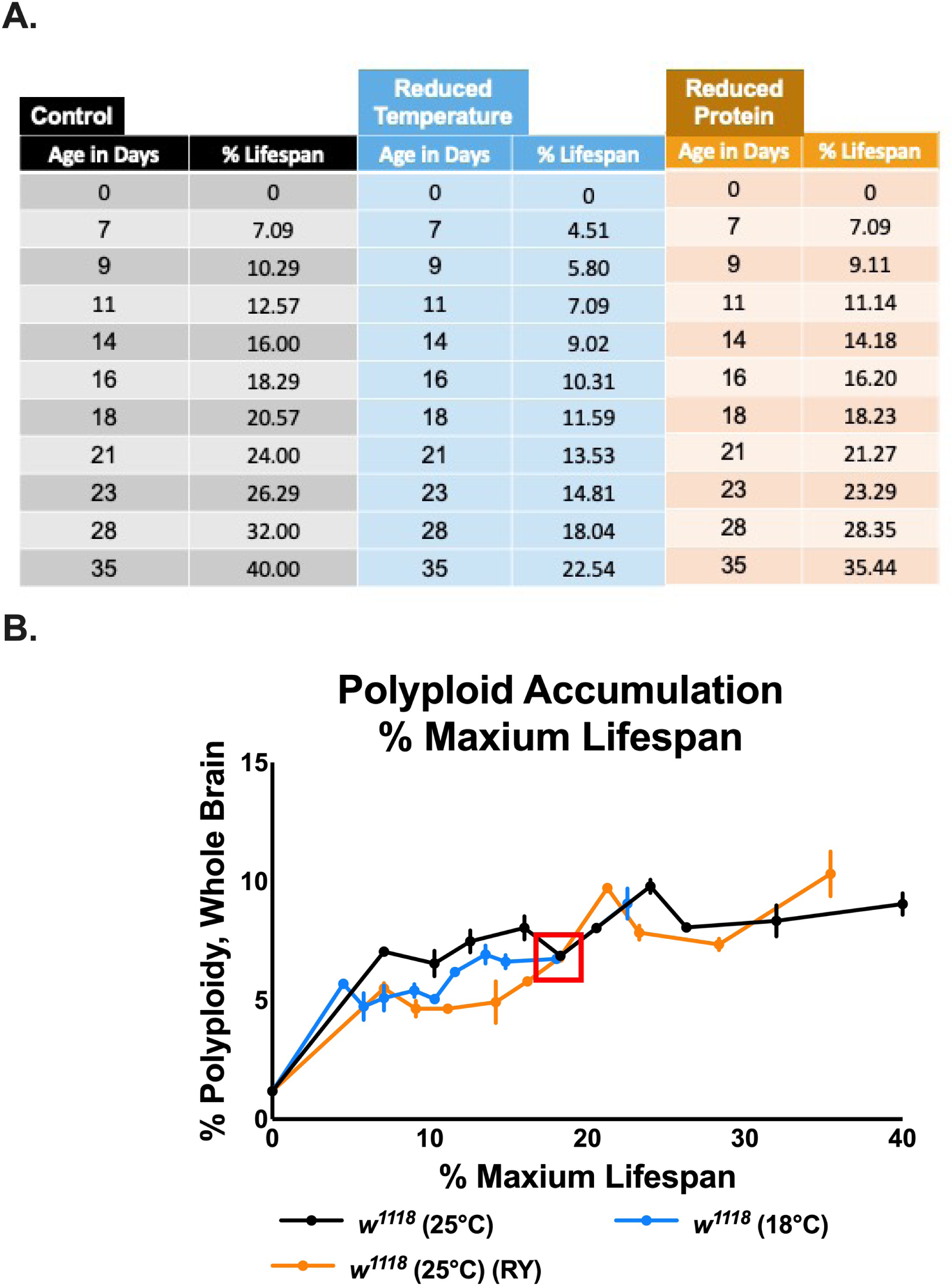
The cell cycle re-entry window for cells in the adult fly brain is flexible and impacted by temperature and diet. (A.) A table displaying chronological age with the percent of the maximum lifespan for conditions that prolong lifespan. (B.) The proportion of polyploid cells in the brain vs. percent of the maximum lifespan. Reduced temperature and reduced yeast slow the accumulation of polyploid cells, even when normalized to percent of maximum lifespan. Red box indicate when polyploidy levels under reduced temperature or reduced yeast reach control levels.

A third possibility is a portion of cells re-enter the cell cycle, become polyploid, and then with age the cell death rate of the polyploid cells increases, such that the total level of polyploidy remains constant. Our data is not fully consistent with this model since, under control aging conditions, the proportion of diploid cell death also significantly increases with age (Fig.1E.) and at each timepoint measured is significantly higher than polyploid cell death, which remains relatively constant during aging (Fig.1E).

Alternatively, brain region differences in polyploid cell accumulation and cell death may result in an overall net gain of zero polyploid cells in later adulthood. Cells in different brain regions may be in different states of G0, since the proportion of polyploidy is not the same in the OLs and CB (Fig. 6 & 7). Furthermore, cells in one region of the brain may have a more flexible G0 state, since OLs of flies housed in different conditions accumulate polyploidy slower (Fig. 6 & 7), suggesting their G0 state became more robust under different conditions. Concurrent to altered states of G0 and flexibility in the G0 state, cell death in the CB is higher than the OLs (Fig. 6 &7) and is influenced by different environmental conditions. Taken together, our data supports more complex models where there is an intricate relationship between brain region specific polyploid accumulation and brain region specific cell death, possibly mediated by mTOR signaling. Future studies will be needed to elucidate how these variables all contribute to polyploidy in the aging brain.

**Figure 6:**
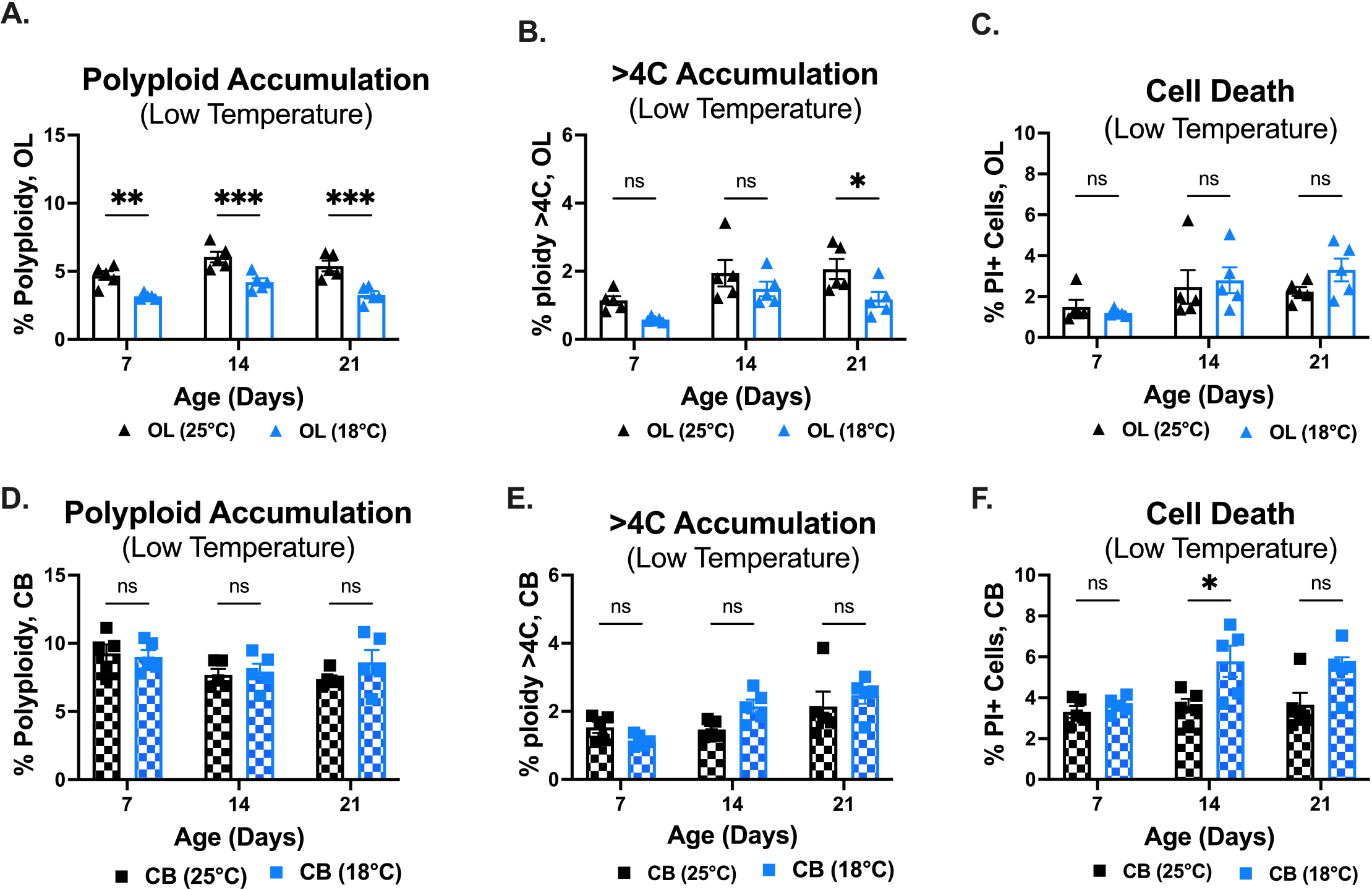
Housing flies at a low temperature impacts polyploidy in the optic lobes more than the central brain. (A.) Flies housed at 18°C accumulate fewer polyploid cells in their optic lobes (OL) (5 biological replicates per time point, 4 optic lobes per biological replicate 2-way ANOVA, Aging factor, p=0.002, Temperature factor, p<0.0001, Sidak’s multiple comparison’s test). (B.) The proportion of cells with more than 4c DNA content is reduced in flies housed at 18°C (5 biological replicates per time point, 4 OL per biological replicate 2-way ANOVA, Aging factor, p=0.0031, Temperature factor, p=0.0037, Sidak’s multiple comparison’s test). (C.) Housing temperature does not significantly impact cell death (5 biological replicates per time point, 4 OLs per biological replicate, 2-way ANOVA, Aging factor, p=0.018). (D.) Temperature does not affect polyploid accumulation in the central brain (CB) (2-way ANOVA, Temperature factor, p=0.420). (E.) The proportion of cells that exceed a 4C DNA content increases with age regardless of housing temperature (2-way ANOVA, Aging factor, p=0.002). (F.) Low temperature impacts cell death in the CB (2-way ANOVA, Aging factor, p=0.042, Temperature factor, p=0.0018). Graphs display individual biological replicates, with SEM. All significant differences are noted: *p<.05, **p<.01 \*\*\**p*<0.001, ***p<0.0001. Abbreviations: Optic Lobe (OL), Central brain (CB)

**Figure 7:**
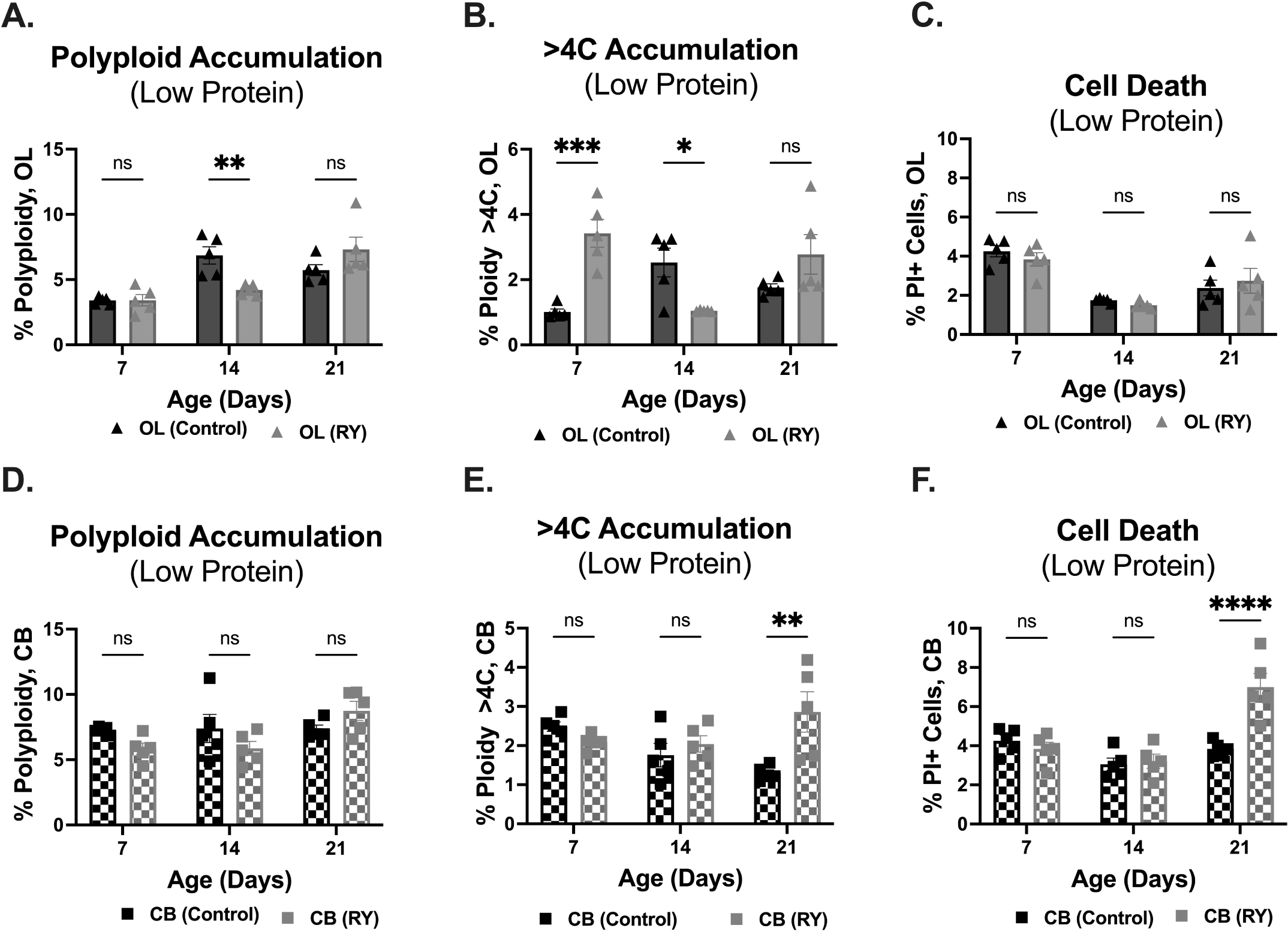
The impact of a low-protein diet on polyploidy is brain region specific. (A.) At age day 14, flies fed a low-protein diet have fewer polyploid cells in their optic lobes (OL) (5 biological replicates per time point, 4 OL per biological replicate, 2-way ANOVA, Aging factor, p<0.0001, Interaction, p=0.0021, Sidak’s multiple comparison’s test). (B.) A low-protein diet influences the number of cells that exceed a 4C DNA content in the OL of flies fed (5 biological replicates per time point, 4 OL per biological replicate, 2-way ANOVA, Aging factor, p=0.347, Interaction, p<0.0001, Sidak’s multiple comparison’s test). (C.) Regardless of diet, cell death in the OL decreased during aging (2-way ANOVA, Aging factor, p<0.0001). (D.) Polyploid accumulation in the central brain (CB) was not influenced by diet (5 biological replicates per time point, 2 CB per biological replicate, 2-way ANOVA, Aging factor, p=0.0394, Interaction, p=0.042, Sidak’s multiple comparison’s test). (E.) At age day 21, flies on a low-protein diet accumulated more cells with a DNA content exceeding 4C in the CB (5 biological replicates per time point, 2 CB per biological replicate, 2-way ANOVA, Diet, p=0.035, Interaction, p=0.0042, Sidak’s p multiple comparison’s test). (F.) A low-protein diet alters cell death in the CB (5 biological replicates per time point, 2 CB per biological replicate, 2-way ANOVA, Aging factor, p<0.0001, Interaction, p=0.0003, Diet factor, p=0.0068, Sidak’s multiple comparison’s test test). Graphs display individual biological replicates, with SEM. All significant differences are noted: *p<.05, **p<.01 \*\*\**p*<0.001, ***p<0.0001. Abbreviations: Optic Lobe (OL), Central brain (CB)

## Acknowledgments and Funding

The authors thank the members of the Buttitta lab and Dr. Scott Pletcher for advice and discussions. This work was supported by the National Institute on Aging Career Training in the Biology of Aging Training Grant (NIH T32 AG000114) and by the National Institute of General Medical Sciences of the National Institutes of Health (NIH R35 GM149273). Stocks obtained from the Bloomington Drosophila Stock Center (NIH P40OD018537) were used in this study. The anti-Lamin (DSHB ALD67.10) antibody developed by Dr. Paul Fisher (SUNY Stony Brook) was obtained from the Developmental Studies Hybridoma Bank, created by the NICHD of the NIH and maintained at The University of Iowa, Department of Biology, Iowa City, IA 52242.

## Authorship

D.D. performed most of the data acquisition, analysis, and manuscription preparation with input from L.B. J.S. and M.D. aided with lifespans, flow cytometry and data analysis.

