## Supplementary Figures for "Cell cycle re-entry in the aging *Drosophila* brain"

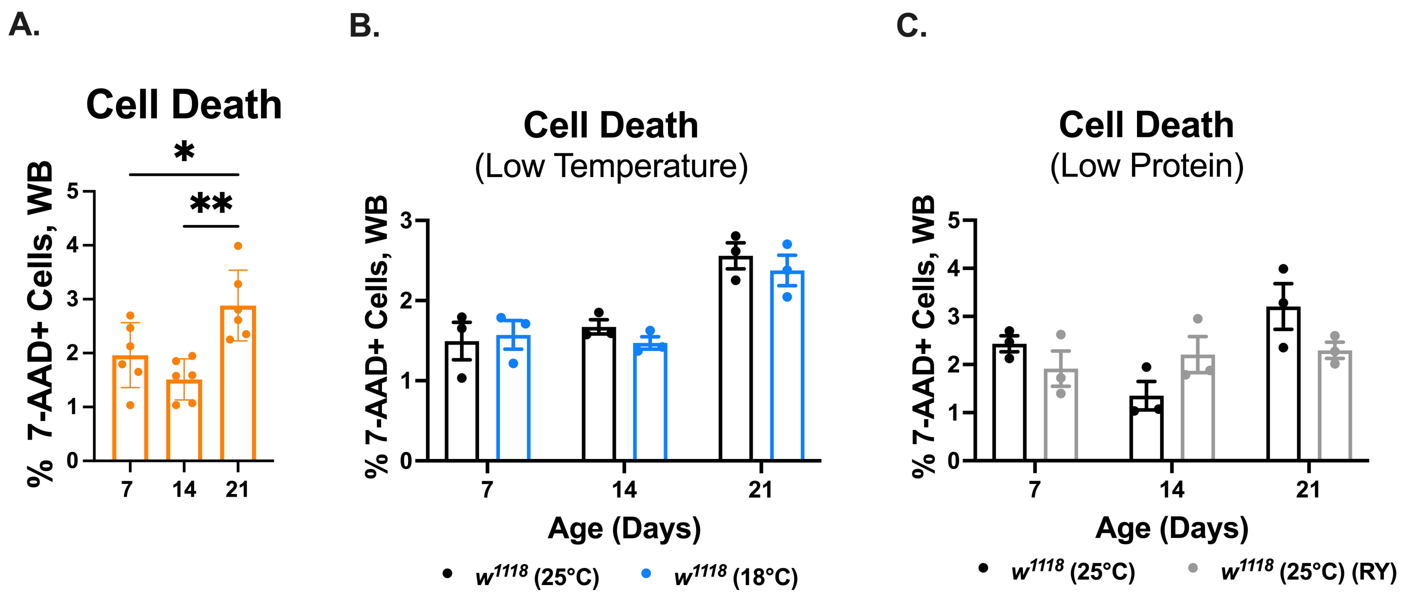


**Supplementary Figure 1: 7-AAD, a cell viability dye, shows similar results as propidium iodide.** (A). Cell death significantly increases during aging in wild type (*w*$^{1118}$), control flies (Ordinary one-way ANOVA, p=0.0023, Tukey's multiple comparisons test). (B.). There is no significant temperature effect on cell death (2-way ANOVA, Temperature effect, p=0.426, Aging effect, p=0.001). (C.) A low protein diet does not significantly influence cell death (2-way ANOVA, Temperature effect, p=0.487, Aging effect, p=0.036). Graphs display individual biological replicates, with SEM. Each biological replicate contains one male and one female brain. All significant differences are noted: *p<.05, **p<.01.

A.

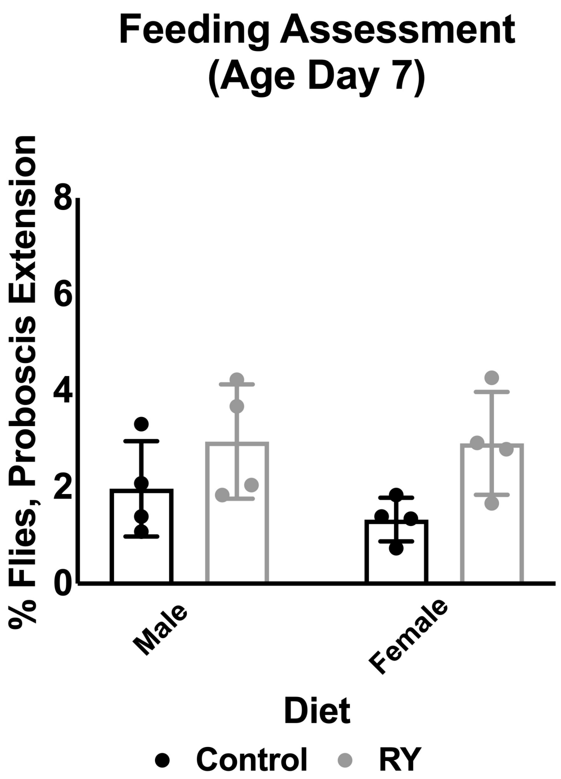


B.

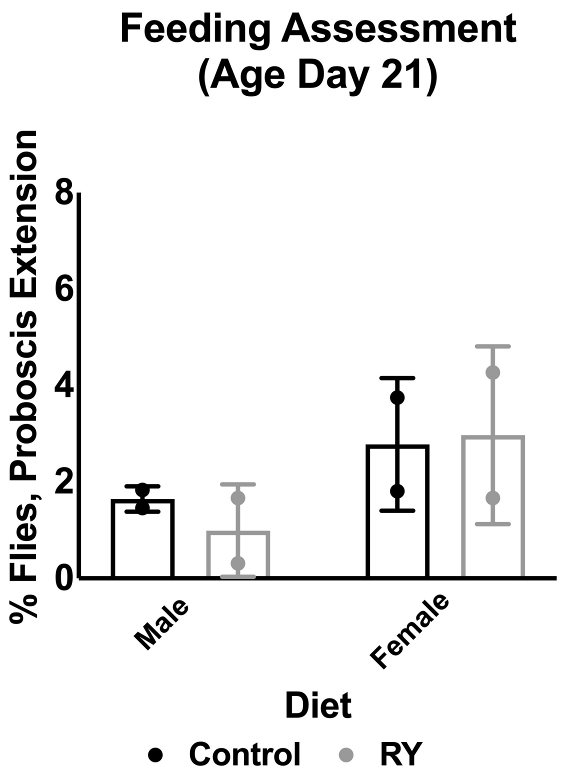


**Supplementary Figure 2: Feeding rate is not different between flies fed a control diet or reduced yeast diet.** (A.). At age day 7, the feeding rate of flies fed a reduced yeast diet is similar to controls (2-way ANOVA, diet factor p=0.02). (B.) At age day 21, the feeding rate of flies on different diets is not different (2-way ANOVA, diet factor, p=0.80).

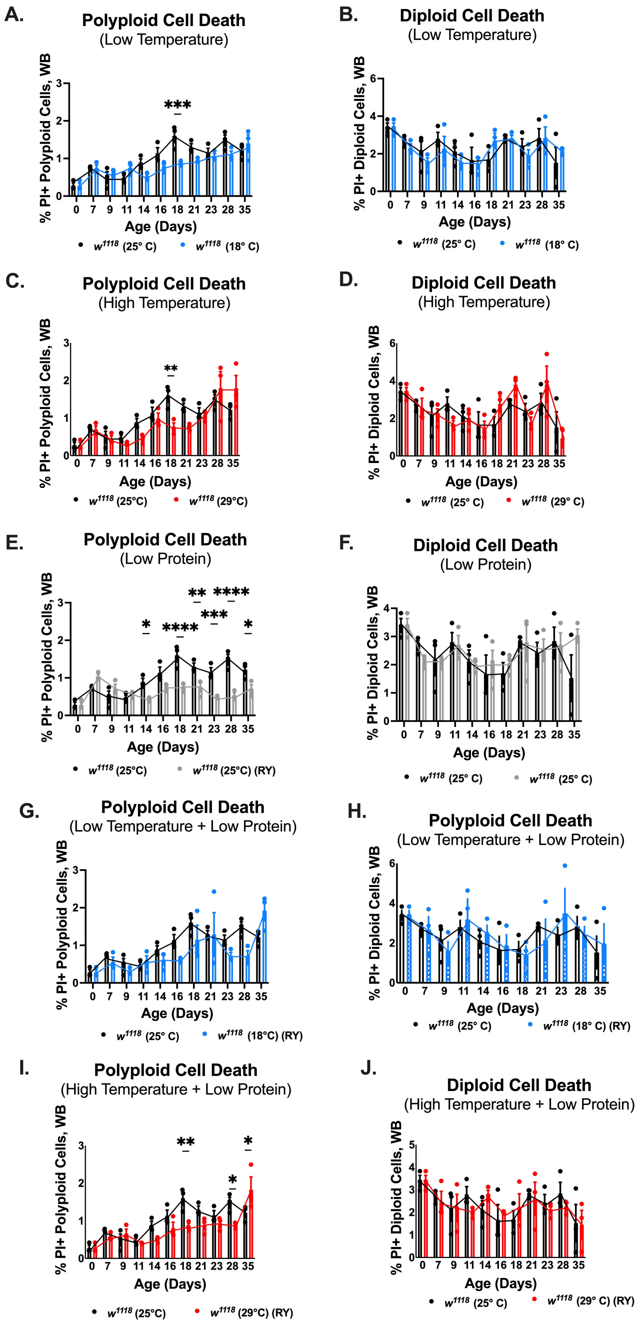

**Supplementary Figure 3: Housing conditions reduce polyploid cell death.** (A.) Housing flies at a low temperature significantly reduces polyploid cell death during aging (2-way ANOVA, Aging effect, p<0.0001, Temperature effect, p=0.0031, interaction, p=0.009, Sidak's multiple comparisons test). (B.) Diploid cell death significantly increase with age but is not significantly affected by low temperature aging (2-way ANOVA, Aging effect, p<0.0017, Temperature effect, p=0.50, Sidak's multiple comparisons test). (C.) Housing flies at a high temperature did not affect polyploid cell death during aging (2-way ANOVA, Temperature effect, p=0.063). (D.) A high housing temperature does not significantly influence diploid cell death (2-way ANOVA, Aging effect, p<0.0001, Temperature effect, p=0.916, Sidak's multiple comparisons test). (E.) Feeding a reduced protein diet significantly reduces polyploid cell death aging (2-way ANOVA, Aging effect, p<0.0001, diet effect, p<0.0001, interaction, p<0.0001, Sidak's multiple comparisons test). (F.) A reduced protein diet does not influence diploid cell death (2-way ANOVA, Aging effect, p=0.016, Diet effect, p=0.517, Sidak's multiple comparisons test). (G.) Housing flies at a low temperature and feeding a reduced protein diet significantly reduces polyploid cell death during aging (2-way ANOVA, Aging effect, p<0.0001, Temperature and diet effect, p=0.0314, Sidak's multiple comparisons test). (H.) Housing flies at a low temperature and feeding a reduced protein diet had no significant effect on diploid cell death (2-way ANOVA, Aging effect, p=0.083, Temperature & Diet effect, p=0.935). (I.) When flies housed at a high temperature and fed a low protein diet, polyploid cell death is significantly lower (2-way ANOVA, Aging effect, p<0.0001, Temperature effect, p=0.001, Interaction, p=0.0003, Sidak's multiple comparisons test). (J.) Housing flies at a high temperature and feeding a low protein diet had no significant effect on diploid cell death (2-way ANOVA, Aging effect, p=0.0174, Temperature & Diet effect, p=0.749). Graphs display individual biological replicates, with SEM. Each biological replicate contains one male and one female brain. All significant differences are noted: *p<.05, **p<.01 ****p*<0.001, ***p<0.0001.
